# Inhibiting CRF Projections from the Central Amygdala to Lateral Hypothalamus and Amygdala Deletion of CRF Alters Binge-Like Ethanol Drinking in a Sex-Dependent Manner

**DOI:** 10.1101/2024.04.09.588750

**Authors:** Sophie C. Bendrath, Hernán G. Méndez, Anne M. Dankert, Jose Manuel Lerma-Cabrera, Francisca Carvajal, Ana Paula Dornellas-Loper, Sophia Lee, Sofia Neira, Harold Haun, Eric Delpire, Montserrat Navarro, Thomas L. Kash, Todd E. Thiele

**Author notes:** Co-First Authors (Sophie C. Bendrath, Hernán G. Méndez). Corresponding Author: Todd E. Thiele, Ph.D., Department of Psychology & Neuroscience University of North Carolina at Chapel Hill Chapel Hill, NC 27599-3270.

## Abstract

**Background:** Binge alcohol drinking is a dangerous pattern of consumption that can contribute to the development of more severe alcohol use disorders (AUDs). Importantly, the rate and severity of AUDs has historically differed between men and women, suggesting that there may be sex differences in the central mechanisms that modulate alcohol (ethanol) consumption. Corticotropin releasing factor (CRF) is a centrally expressed neuropeptide that has been implicated in the modulation of binge-like ethanol intake, and emerging data highlight sex differences in central CRF systems.

**Methods:** In the present report we characterized CRF+ neurocircuitry arising from the central nucleus of the amygdala (CeA) and innervating the lateral hypothalamus (LH) in the modulation of binge-like ethanol intake in male and female mice.

**Results:** Using chemogenetic tools we found that silencing the CRF+ CeA to LH circuit significantly blunted binge-like ethanol intake in male, but not female, mice. Consistently, genetic deletion of CRF from neurons of the CeA blunted ethanol intake exclusively in male mice. Furthermore, pharmacological blockade of the CRF type-1 receptor (CRF1R) in the LH significantly reduced binge-like ethanol intake in male mice only, while CRF2R activation in the LH failed to alter ethanol intake in either sex. Finally, a history of binge-like ethanol drinking blunted CRF mRNA in the CeA regardless of sex.

**Conclusions:** These observations provide novel evidence that CRF+ CeA to LH neurocircuitry modulates binge-like ethanol intake in male, but not female mice, which may provide insight into the mechanisms that guide known sex differences in binge-like ethanol intake.

## INTRODUCTION

In the United States, alcohol (ethanol) binge drinking is a public health concern not only associated with many negative acute and long-term neuropsychological effects, but also with the development of alcohol use disorders (AUDs) ^1–5^. The National Institute on Alcohol Abuse and Alcohol (NIAAA) defines binge drinking as alcohol consumed within a 2-hour timeframe that leads to blood ethanol concentrations (BECs) of 80mg/dL or greater^6^. Males have been more prominently diagnosed with AUD’s^7^, however, the numbers of female AUD diagnoses has recently risen^7,8^. Interestingly, there seem to be distinct differences between the male and female drinking populations, such that a history of trauma and acute life stressors influences alcohol craving and relapse almost exclusively in females as compared to males^9,10^. This suggests an underlying common circuitry between stress and alcohol craving that may be sex-specific, yet remains largely unknown.

Corticotropin releasing factor (CRF) is a pro-stress peptide in the extended amygdala that becomes dysregulated during alcohol use and addiction^11^. The central nucleus of the amygdala (CeA) is a particularly important region in relation to AUD, integrating stress and reward reactions to events to form behavioral responses^12^. Previous work from our lab using “drinking in the dark” (DID) procedures to model binge-like ethanol intake in male mice has shown that 1 and 3, 4-day cycles of binge-like ethanol intake promote increased CRF expression in the central amygdala (CeA)^13^ as well as long-lasting subsequent increases of voluntary ethanol intake^14^, suggesting extended plasticity stemming from a history of binge-like ethanol intake. During acute alcohol administration in rats, the CeA exhibits augmented release of the CRF peptide^15^ which persists into withdrawal, and infusion of CRF into the CeA increases anxiety-like behavior during abstinence, suggesting that CRF may play a role in alcohol craving^16,17^. Additionally, the CeA is the main output area related to behavioral responses to stress, and has strong projections to other pro-stress areas, such as the lateral hypothalamus (LH)^11,18^. However, the role of CRF projections from the CeA to the LH in modulating binge-like ethanol consumption has not been examined.

The LH is a heterogeneous brain area expressing a variety of neuropeptide systems and has been linked to a number of brain regions that modulate stress and reward^18^. Stress-induced avoidance behavior in male rats is directly modulated by a neuronal connection from the CeA to the LH, functionally linking these two brain areas^19^. Moreover, CRF in the LH is dysregulated during both chronic alcohol use and withdrawal^20^, and CRF mRNA levels are elevated in the hypothalamus after acute alcohol administration^21^. On the receptor level, LH CRF1 receptor (CRF1R) and CRF2 receptor (CRF2R) mRNA expression and ethanol consumption are positively correlated in male rats^22^, and CRF1R in the LH modulate stress responses^23^, thus CRF1R and CRF2R in the LH may be potential targets for CRF projections from the CeA that modulate binge-like ethanol drinking.

Importantly, the role of CeA CRF+ neuronal innervation of the LH and CRF receptor signaling in the LH in the modulation of binge-like ethanol intake, as well as potential sex differences, are not well understood. Some evidence suggests that female rats are more likely to self-stimulate the lateral hypothalamus than males^24^, making this a sex-specific hedonic brain area. Additionally, female rodents are predisposed to anxiety disorders and higher ethanol consumption than males, which is largely influenced by differential CRF signaling in the extended amygdala^25^. Female mice lacking beta endorphin had higher ethanol consumption and CRF mRNA than males in the extended amygdala^26^. Additionally, relative to males CRF receptor binding is increased in the cortex and amygdala of female rats^25^ and CRF mRNA expression is greater in the amygdala of female mice^27^. At the level of cell signaling, acute application of ethanol onto mouse CeA slice preparations reduced GABA release onto CRF1R-expressing neurons in male, but not female mice, and application of exogenous CRF increased the firing rate of CRF1R-expressing neurons to a greater extent in male mice^28^. Thus, while there is emerging evidence of sex differences in amygdala and hypothalamic CRF systems and how they respond to ethanol, much more work is necessary to further unpack the mechanisms that are involved.

In light of the converging evidence that CRF signaling in the CeA and LH modulate ethanol intake, and evidence of a CRF+ neurocircuit arising from the CeA and innervating the LH, the current study was aimed at assessing the role of a CRF+ pathway from the CeA to the LH and CRF receptor signaling in the LH in the modulation of binge-like ethanol drinking. Additionally, experiments were conducted to investigate whether binge-like ethanol consumption alters CRF and CRF receptor mRNA in these brain regions, and if genetic deletion of CRF or CRF1R in the neurons of the CeA would impact binge-like ethanol intake. Importantly, because there has been very little investigation into potential sex differences in the mechanisms by which CRF signaling modulates binge-like ethanol intake, *in vivo* pharmacology, genetic deletion, chemogenetic studies, and quantitative polymerase chain reaction (qPCR) experiments described below entailed both male and female mice.

## METHODS

### Animals

Male and female C57BL/6J mice (Jackson Laboratories, Bar Harbor, ME) aged 8-10 weeks at the start of experimentation were used for the pharmacological study (CRF1-antagonist: male, n=20; female n=20) and the qPCR study (male, n=25; female, n=25). The *in vivo* chemogenetic experiments used 30 male and female (Cre+ male, n=7; Cre+ female, n=8; Cre-male, n=8; Cre-female, n=7) CRH-ires-Cre (CRH-Cre) mice (positive for the expression of Cre under the CRH promoter as determined by standard PCR genotyping protocols) at least 10 weeks of age. CRH-Cre mice were generated and genotyped as described previously^29^. For the Crh deletion experiment, 27 CRF floxed males (Control treated male, n=14; Cre treated male, n=13) and 26 CRF floxed females (Control treated female, n=13; Cre treated female, n=14) aged 11-13 weeks at the start of behavioral testing were used with mice receiving either infusion of an AAV8-hsyn-gfp (Control) or AAV8-hsyn-cre-gfp (Cre) into the CeA as described below. Floxed CRF mice were generated as described previously^30^. For the Crhr1 deletion experiment, 31 CRFR1 floxed male (Control treated male, n=14; Cre treated male, n=17) and 20 CRF1R floxed females (Control treated female, n=8; Cre treated female, n=12) aged 15-22 weeks at the start of behavioral testing were used with mice receiving either infusion of an AAV8-hsyn-cre-gfp (Cre) or a AAV8-hsyn-GFP (Control) into the CeA as described below. CRF1R floxed mice were generated as described in the supplementary information extended methods. All mice were housed individually in an AAALAC accredited vivarium on a reverse 12 h light-dark cycle, with the lights going off at 09:30 or 10:30 hours, depending on the room. Water (unless otherwise stated) and food (Prolab® RMH 3000 (Purina LabDiet®; St. Louis, MO)) were available to all animals ad libitum. All procedures were approved by the University of North Carolina Institutional Animal Care and Use Committee, and conducted in accordance with the Guidelines for the Care and Use of Laboratory Animals.

### DREADD and Cannula Surgery

Surgeries were conducted on an Angle II™ Stereotax (Leica Instruments, Buffalo Grove, IL). Animals received intraperitoneal (i.p.) injections (1.5 mL/kg) of ketamine (117 mg/kg)/ xylazine (7.92 mg/kg), with all coordinates being measured from bregma. For the *in vivo* chemogenetic study, mice received either an injection of Cre-dependent control vector (AAV8-hSyn-DIO-mCherry, Addgene, Watertown, MA) or the Cre-dependent Gi/o-coupled DREADD vector (AAV8-hSyn-DIO-hM4d-mCherry, Addgene, Watertown, MA) into the CeA (AP −1.06, ML +/-2.42, DV −4.63). The injection needle remained in place for 10-15 additional minutes before being withdrawn. This Designer Receptor Exclusive Activated by Designer Drugs (DREADD) has been validated and used to manipulate CRF activity in the central amygdala previously^31–33^. Additionally, bilateral guide cannulas (Plastics One; Roanoke, VA) were implanted into the LH (AP −1.10, ML +/-1.10, DV −5.10; injector: 5MM with 0.5MM projection). Upon surgery completion, all mice recovered for at least 3 weeks before starting behavioral testing for maximal viral expression. For the *in vivo* pharmacological study, animals were implanted with bilateral guide cannulas in the same LH coordinates for intra-LH microinfusions, and were allowed to recover from surgery for 1 week before starting the DID procedure.

### Genetic Deletion Surgery

Surgeries were conducted on Kopf Stereotaxic device (Kopf Instruments, Tujunga, CA, USA). Mice were first anesthetized by placing inside a chamber that contained vaporized 4% isoflurane and oxygen and then maintained via at 2-3% isoflurane during surgery. All coordinates being measured from bregma. A Hamilton syringe infused 250nl of either control virus (AAV8-hsyn-GFP) and titer of 4.1X10e12 or Cre virus (AAV8-hsyn-Cre-GFP) titer of 4.5X10e12 to the CeA at (−0.62, ML +/-2.70, DV −5.00 or −0.80, ML +/-2.95, DV −4.85). The injection needle remained in place for approximately 7 additional minutes per injection site before being withdrawn. Upon surgery completion, all mice recovered for at least 3 to 6 weeks before starting behavioral testing for maximal viral expression. For the study we included mice with CeA bilateral and unilateral hits, a hit was defined as having viral tag GFP presence in the CeA.

### “Drinking in the Dark” (DID) Procedures

A 4-day DID model was used to assess ethanol intake, which is a standard protocol to induce binge-like ethanol consumption^34^. Please refer to extended methods on supplemental materials for full methodology.

### Drug Administration

Approximately 30 minutes prior to ethanol access on the 4th day of DID, animals received microinjections with either the DREADD ligand clozapine-N-oxide (CNO; 900pmol) or vehicle (1% DMSO in 0.9% saline), in a counterbalanced 2×2 Latin-square design during consecutive weekly DID session. All animals were assigned randomly to either drug or vehicle treatment on the first test day, then received the other treatment on test day 2 during the second DID cycle. Drug injections were performed with a Hamilton syringe (Reno, NV) on a Harvard Apparatus PHD 2000 infusion pump (Holliston, MS) at a rate of 0.10µl for 3 minutes (0.30µl total). After injection, infusion needles stayed in the cannula for an additional minute to ensure complete diffusion. Tail blood samples were collected by nicking the lateral tail vein on the 4th day immediately after ethanol access (about 30µl), to assess blood ethanol concentrations (BECs) on an (AM1) Alcohol Analyzer (Analox, London, UK).

### In Vivo Pharmacology

For CRF1R antagonism, NBI-35965 hydrochloride (Tocris, Bristol, UK: 30pmol/0.3 µl/side) was dissolved in sterile H_2_O for a drug make-up of 13µg/ml. Bilateral microinjections of NBI or vehicle occurred about 30 minutes before test day ethanol access (DID day 4), and were administered at a rate of 0.10µl/min to reach the target volume of 0.30µl per side using Hamilton infusion pumps. For CRF2R agonism, Urocortin 3 (Ucn3; GenScript USA, Piscataway, NJ: 60pmol/0.4μl/side) was dissolved in DMSO (10% v/v final concentration; Sigma-Aldrich, St. Louis, MO) and diluted with 0.9% saline, for a target concentration of 250µg/mL. Ucn3 or its vehicle was also administered at a rate of 0.1μl/min 30 minutes before ethanol access. Injectors were left in place for one minute post-infusion to allow for diffusion of the drug away from the injector tip and minimize back flow of drug as the injectors were removed. Again, all animals were assigned randomly to either the drug or vehicle groups, and received the opposite treatment on test days in a 2×2 Latin square design during consecutive weekly DID session.

### Perfusion, Histology, Fluorescence In Situ Hybridization (FISH), and Reverse Transcription Quantitative Polymerase Chain Reaction (RT-qPCR)

In order to confirm genetic deletion of CRF and CRF1R, brain collection, cryosectioning and FISH was performed as previously described^35^. RNA extraction, cDNA synthesis, and RT-qPCR were performed as previously described^36,37^. Please refer to extended methods on supplemental materials for full methodology.

### Statistical Analysis

All analyses and graphs were generated with SPSS (IBM Analytics, Armonk, New York) and GraphPad Prism (GraphPad Software, Inc. La Jolla, CA, USA). Unpaired two-tailed t-tests, or repeated measures analysis of variance (ANOVA) were used where appropriate, to assess experimental treatment effects on ethanol intake, BEC’s and sucrose intake. Relative expression of mRNA was analyzed using the comparative CT (ΔΔCT) method. Two-way ANOVAs, or Kruskal-Wallis tests for non-parametric data, were performed to examine the effects of repeated cycles of DID of mRNA expression. Bonferroni corrected t-tests and planned comparisons were used for significant ANOVA effects where appropriate. All data are reported as the mean ± standard error of the mean considered significant at p < 0.05 (two-tailed). For our genetic deletion experiments we used Repeated Measures Two Way ANOVA, student’s t-test and Mixed effect analysis. If main effects were present, Šídák multiple comparison post hoc test was performed, p value smaller than 0.05 was considered significant. Both unilateral and bilateral hits were included in this study. Data points were removed according to exclusion criteria when applicable, see extended methods in supplemental.

## RESULTS

### Gi DREADD silencing of the CeA to LH pathways revealed a sex-specific reduction for ethanol drinking in males only

Placing the *Gi* DREADD into the CeA, and activating it with CNO injections from the LH, there was a trend towards significance with treatment (F(1,26)=3.76, p=0.063), and a significant effect of sex (F(1,26)=17.68, p< 0.001) on ethanol intake. The interaction between sex and treatment also had a significant impact on ethanol consumption (F(1,26)= 4.86, p=0.04). Planned comparisons for vehicle and CNO injection groups of each sex showed that CNO injections significantly reduce binge-like ethanol drinking in males (p=0.009), but not females (p=0.861) (***Fig 1A***). Blood ethanol concentrations (BECs) showed a significant main effect for injection type (F(1,26)= 4.114, p=0.053), as well as sex (F(1,26)= 8.19, p=0.008). The interaction of sex and treatment was not a significant factor of BECs (F(1,26)=2.444, p=0.130). Again, this effect reflects a significant reduction in BEC’s for males injected with CNO relative to vehicle (p=0.021), and not females (p=0.736) (***Fig 1B***). For DID sucrose drinking there was no significant impact of treatment (F(1,26)=3.545, p=0,071). The main effect of sex on sucrose consumption was significant, such that females overall drank more than males (F(1,26)= 7.47, p=0.01). The interaction of sex and treatment again did not significantly affect sucrose intake (F(1,26)=0.320, p=0.576). (***Fig 1C***). **Fig 1D** shows a representative image of GI DREADD expression in the CeA, and **Fig 1E** shows the cannulae placement map.

**Fig. 1.**
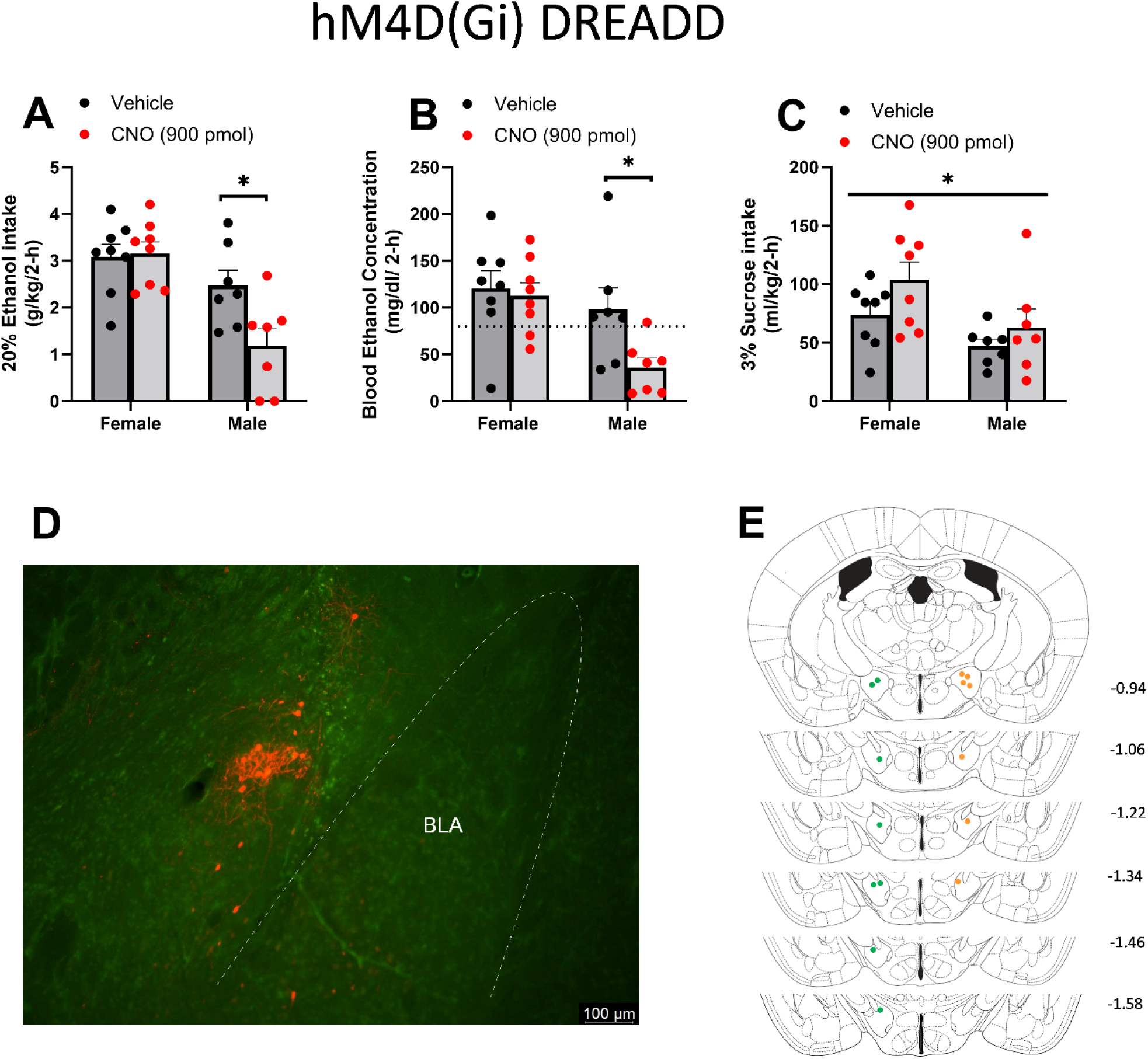
Chemogenetic silencing the the CRF+ CeA → LH circuit. (A) Only males show a significant reduction in binge-like ethanol intake when the CRF+ CeA-LH circuitry is inhibited. (B) BEC’s show a similar reduction during CNO treatment in males only (dotted line indicates 80mg/dL). (C) No sex-specific blunting is seen in females or males for sucrose intake. (D) Representative photomicrograph of Gi/o DREADD expression in the CeA. (E) Cannula placement for Gi DREADD animals; placements are marked to the left (green) for females, and right (orange) for males. Due to the cannula being placed within a pedestal, only one hemisphere is shown for placement as each subject was consistent on both hemispheres. Data are represented as Mean +/-SEM. *p < 0.05.

An *mCherry* control DREADD study was performed in CRH+-cre mice to validate specificity of *Gi* DREADD findings in the DID model. There was no change in ethanol consumption based on sex (F(1,12)=1.921, p=0.191), compound injected (F(1,12)=0.03998, p=0.8449), or an interaction of these two factors (F(1,12)=0.3613, p=0.559) as seen in **Fig 2A**. Similarly, variation in BECs was not a consequence of sex (F(1,12)=0.03295, p=0.859), CNO/Veh injections (F(1,12)=1.142, p=0.3062), or an interplay of the two (F(1,12)=1.33, p=0.2713) as seen in **Fig 2B**. Lastly, when assessing sucrose consumption no significant changes were seen when considering the main effect of sex (F(1,12)=2.488, p=0.1407), treatment (F(1,12)=0.3251, p=0.5791), or an interaction of sex by treatment (F(1,12)=0.01794, p=0.8957), shown in **Fig 2C**. Thus, any changes in ethanol consumption or BECs seen in Gi DREADD animals did not occur in animals using an *mCherry* control DREADD. **Fig 2D** shows a representative image of control DREADD expression in the CeA, and **Fig 2E** shows the cannulae placement map.

**Fig. 2.**
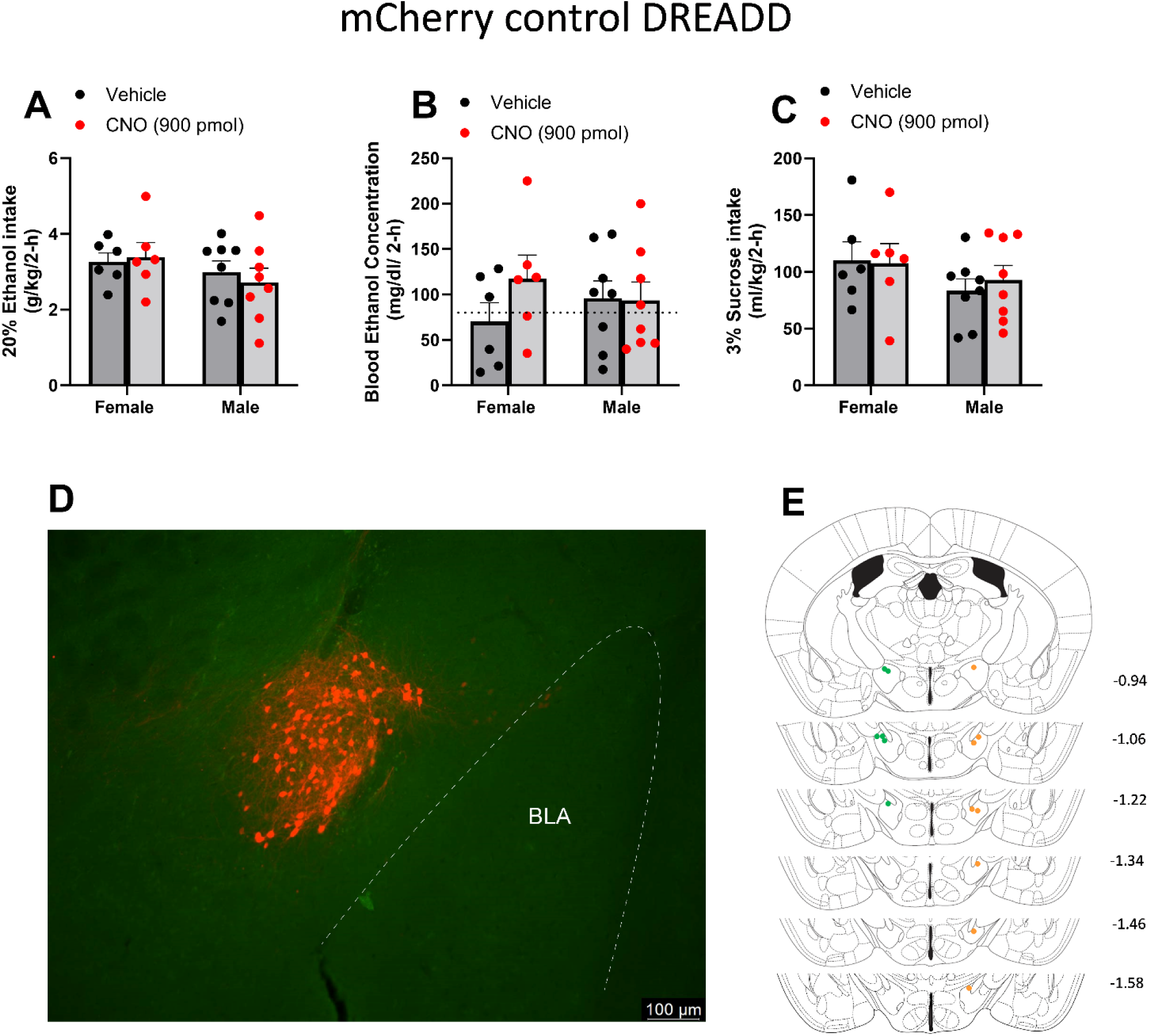
Chemogenetic control experiment. (A) No significant changes are seen in ethanol consumption, (B) BEC’s (dotted line indicates 80mg/dL) (C) or sucrose intake in male and female mice depending on vehicle or CNO treatment. (D) Representative photomicrograph of mCherry control expression in the CeA. (E) Cannula placement for mCherry control animals; placements are marked to the left (green) for females, and right (orange) for males. Due to the cannula being placed within a pedestal, only one hemisphere is shown for placement as each subject was consistent on both hemispheres. Data are represented as Mean +/-SEM. *p < 0.05.

### Pharmacological inhibition of CRF1R in the LH blunts binge-like ethanol intake in male, but not female mice, with no such effects seen during pharmacological activation of CRF2R

To focus on the CRF circuitry within the LH, mice were injected with a CRF1R antagonist (NBI-35965) or CRF2R agonist (Ucn3) into the LH. Relative to vehicle treatment, inhibiting CRF1 receptors in the LH resulted in reduced binge-like ethanol consumption such that the interaction between sex and treatment was significant (F(1,18)= 6.870, p=0.0173), while the main effects of sex (F(1,18)=0.0546, p=0.8179) and treatment (F(1,18)=0.001, p=0.9745) had no significant effect on ethanol consumption. Using paired samples t-test as planned comparisons to assess differences between vehicle and NBI-35965 treatments, there was a significant reduction in ethanol drinking for males with NBI-35965 (t(9)= 2.577, p=0.03), but not in females (t(9)= 1.510, p=0.165) (***Fig 3A***). BEC’s reflect the ethanol drinking data in that only the interaction of sex by compound injection was significant (F(1,35)= 4.854, p=0.0343), while the main effect of sex (F(1,35)=0.4033, p=0.5295) and treatment condition (F(1,35)=2.079, p=0.1582) was not. Planned comparisons again show that the significant CRF1R antagonist-induced reduction in BEC’s occurs in males (p=0.0263), and not females (p>0.9999) (***Fig 3B***). Sucrose drinking remained stable with no significant main effects of sex (F(1,18)=0.4619, p=0.5054), treatment (F(1,18)=0.4645, p=0.5042), or an interaction of these two factors (F(1,18)=0.1778, p=0.6783) (***Fig 3C***). **Fig. 3D** shows the cannulae placement map for the pharmacology experiment. This further supports that, specifically in males, CRF1R’s in the LH are involved in blunting binge-like ethanol drinking.

**Fig. 3.**
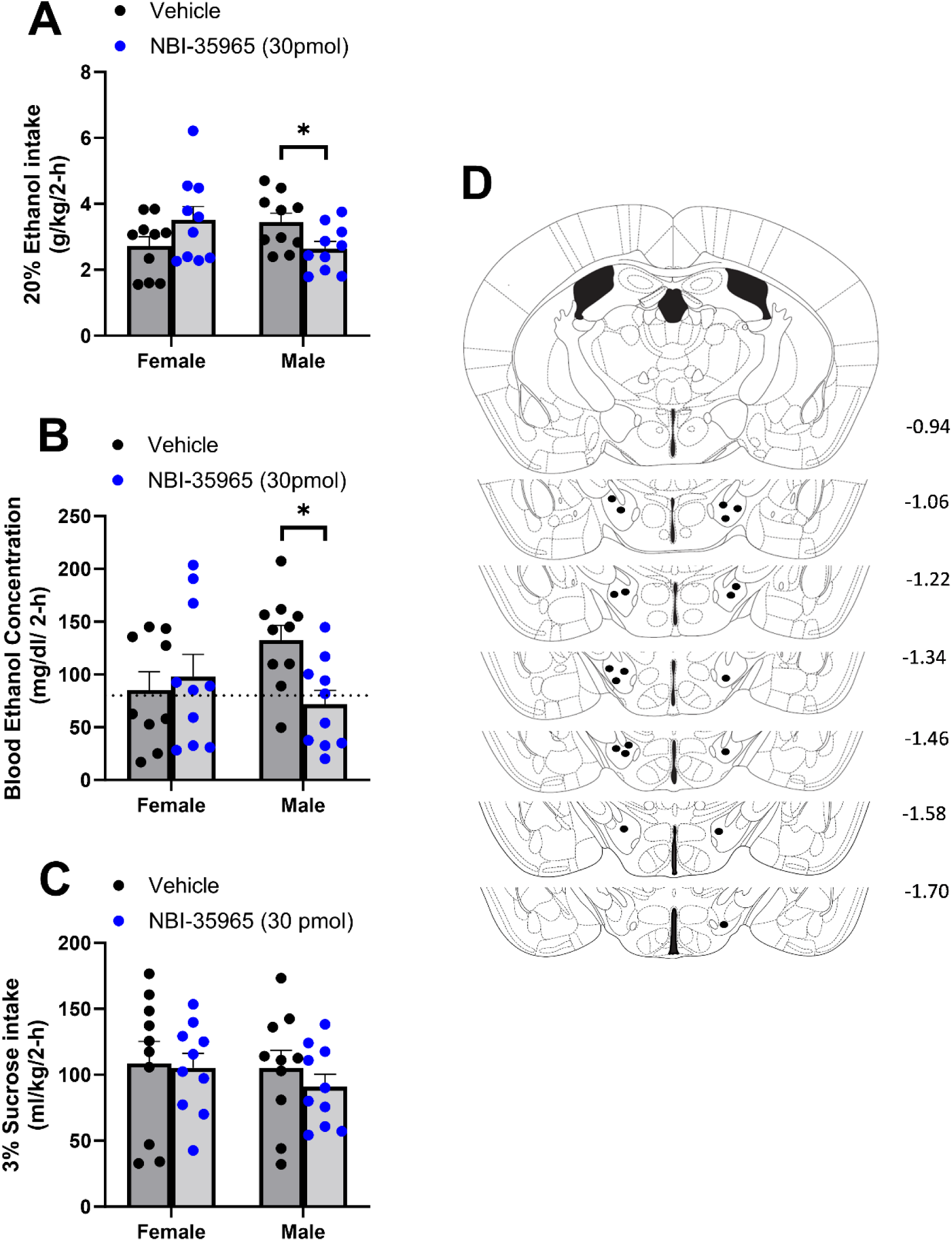
Pharmacological blockade of the CRF1R. (A) Ethanol intake was significantly reduced in males treated with NBI-35965, with no such changes in female ethanol consumption. (B) BECs of male mice were significantly blunted, while female BEC levels remained the same (dotted line indicates 80mg/dL). (C) No significant changes in sucrose consumption between the sexes were observed. (D) Bilateral cannula placements in the LH of all animals. Data are represented as Mean +/-SEM. *p < 0.05.

Next, involvement of the CRF2R was tested via the use of the selective agonist UCN3. Overall, females consumed more ethanol than males (F(1,13)=24.78, p=0.0003), but this effect was independent of whether animals received a UCN3 or vehicle microinjection (F(1,13)=0.6456, p=0.4361). Similarly, there was no significant interaction effect on ethanol consumption when considering sex and compound injected (F(1,13)=0.9005, p=0.3599) (**Fig 4A**). BEC values reflected ethanol binge-drinking findings, such that overall females had higher BECs than males (F(1,13)=35.89, p<0.0001), while drug treatment had no overall effect on BECs (F(1,13)=0.06185, p=0.8075). An interaction of sex x treatment did also not significantly alter BECs (F(1,13)=0.1503, p=0.7045) (**Fig 4B**). Lastly, sucrose consumption did not vary based on sex (F(1,13)=0.8839, p=0.3643), or drug treatment (F(1,13)=0.06682, p=0.8001). Likewise the interaction of sex and drug treatment did not lead to significant changes in sucrose consumption (F(1,13)=0.6334, p=0.4404) (**Fig 4C**). **Fig. 4D** shows the cannula placement for animals included in this pharmacological experiment. Thus, CRF2R agonism in the LH does not appear to significantly alter binge-like ethanol consumption in either male or female mice.

**Fig. 4.**
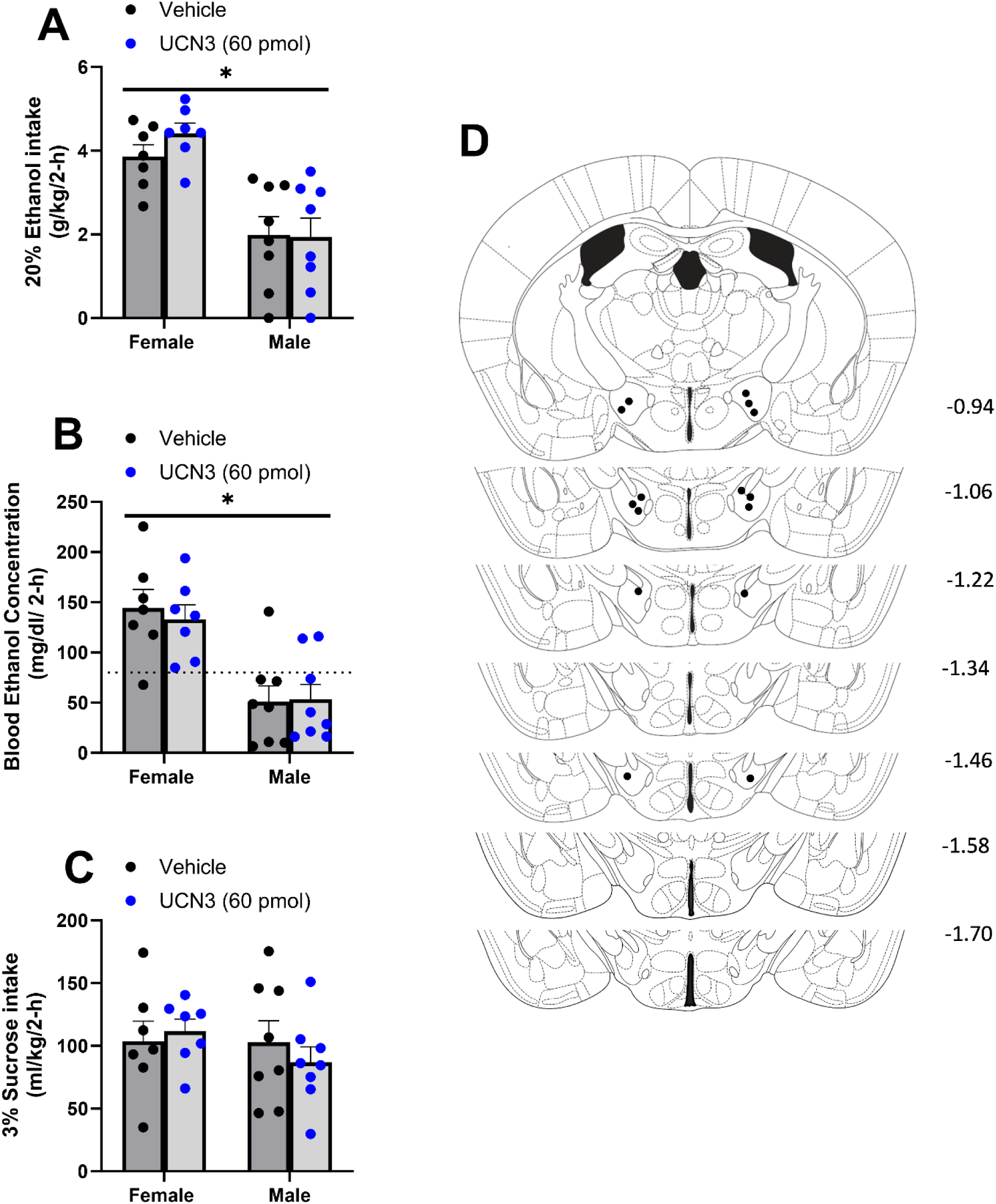
Pharmacological activation of the CRF2R. (A) No changes in ethanol consumption were observed based on UCN3 or vehicle treatment in either sex. (B) Overall females had higher BECs than males, but this effect was independent of UCN3 or vehicle treatment (dotted line indicates 80mg/dL). (C) Sucrose drinking was not affected by Ucn3 injections in a sex-specific manner. (D) Bilateral cannula placements in the LH all animals. Data are represented as Mean +/-SEM. *p < 0.05.

### Repeated cycles of binge-like ethanol consumption alter CRF and CRF receptor mRNA in the amygdala, but not in the LH

Ethanol consumption and BECs associated with the mRNA study are shown in **Fig. 5A and B**, respectively. All groups consumed equal amounts of ethanol during the final week of DID (F(2,32)=1.079, p=0.352), and no group differences in BECs following the final session of DID were observed (F(1,20)=0.332, p=0.571), ensuring that differences in mRNA expression are the result of the number of DID cycles received. A significant main effect of sex revealed that female mice consumed more ethanol than males during the final week of DID (F(1,32)=16.605, p<0.001).

**Fig. 5.**
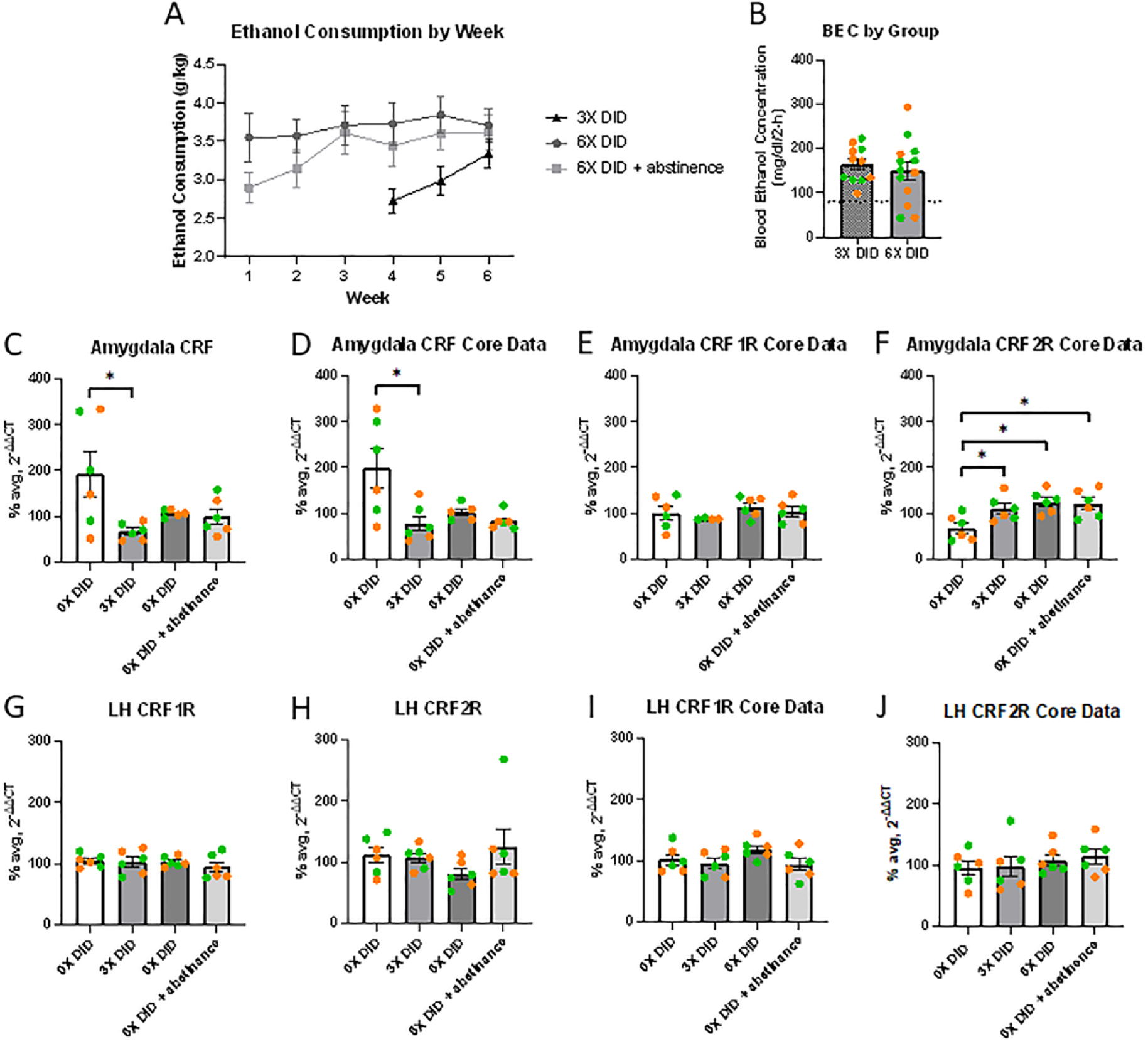
Effects of repeated cycles of binge-like ethanol intake on CRF, CRF1R, and CRF2R mRNA. (A) No group differences in ethanol consumption, (B) or BEC’s (dotted line indicates 80mg/dL) were observed during the final week of DID. (C) CRF mRNA expression in the amygdala was significantly reduced following three cycles of DID, (D) and the same effect was observed by the UNC Advanced Analytics Core. (E) No group differences in CRF1R expression in the amygdala were seen. (F) Mice that received repeated cycles of DID with and without a 24-hour period of abstinence demonstrated greater CRF2R mRNA expression in the amygdala than water control mice. (G) No significant group differences were observed for CRF1R mRNA expression in the LH, (H) or CRF2R mRNA expression in the LH, (I-J) an effect which was replicated by the UNC Advanced Analytics Core. Data are represented as Mean +/-SEM. Female data are indicated by green data points; male data are indicated by orange data points. *p < 0.05.

qPCR was performed to determine the effects of repeated cycles of binge-like ethanol consumption on CRF and CRF receptor mRNA in the amygdala (**Fig. 5C-F**). A two-way ANOVA revealed that CRF mRNA expression in the amygdala was significantly different between groups (F(3,15)=3.313, p=0.049), where mice that received 3 cycles of DID had significantly less CRF mRNA compared to water control mice (**Fig. 5C** p=0.042). No significant main effect of sex or group by sex interaction was found (F(1,15)=0.734, p=0.405; F(3,15)=0.114, p=0.951). As seen in **Fig. 5D**, the same effects of binge-like ethanol consumption on CRF mRNA in the amygdala were observed from the qPCR data generated by the Advanced Analytics Core (Group: F(3,15)=4.377, p=0.021; Sex: F(1,15)=0.519, p=0.482; Group x sex: F(3,15)=0.057, p=0.981). Additional data (**Fig. 5E-F**) from the Advanced Analytics Core demonstrated that there was no main effect or group, sex, or group by sex interaction on CRF1R mRNA expression in the amygdala (F(3,14)=0.597, p=0.628; F(1,14)=0.137, p=0.717; F(3,14)=0.152, p=0.926), but there was an effect of binge-like ethanol consumption on CRF2R mRNA expression in the amygdala. A two-way ANOVA revealed a significant main effect of group (F(3,16)=6.336, p=0.005), and planned comparisons revealed that mice that received three cycles of DID, six cycles of DID, or six cycles of DID followed by a 24 hour period of abstinence showed greater CRF2R mRNA expression in the amygdala compared to water control mice (p=0.049, p=0.009, p=0.014, respectively). No main effect of sex or group by sex interaction was observed (F(1,16)=0.002, p=0.969; F(3,16)=0.793, p=0.516).

qPCR was also performed to determine the effects of repeated cycles of binge-like ethanol consumption on CRF receptor mRNA in the LH (**Fig. 5G-J**). We observed no main effect of group, sex, or group by sex interaction on either CRF1R or CRF2R mRNA expression in the LH (CRF1R: F(3,16)=0.669, p=0.583; F(1,16)=0.350, p=0.562; F(3,16)=1.264, p=0.320; CRF2R: F(3,16)=1.766, p=0.194; F(1,16)=0.377, p=0.548; F(3,16)=1.225, p=0.333). Additionally, no differences were observed in CRF1R or CRF2R mRNA expression in the LH when examining qPCR data produced by the Advanced Analytics Core (CRF1R: F(3,16)=1.645, p=0.219; F(1,16)=0.198, p=0.662; F(3,16)=0.512, p=0.680; CRF2R: F(3,16)=0.668, p=0.584; F(1,16)=0.510, p=0.485; F(3,16)=1.334, p=0.298).

### CeA deletion of CRF

To determine if CRF produced in the CeA plays a role in alcohol drinking, we knocked down CRF in the CeA and measured alcohol consumption in male and female mice using the DID paradigm (**Fig. 6A**). To accomplish this, AAV delivery of Cre in combination with a CRF floxed mouse line was used. This enables site specific deletion of the Crh gene. FISH was used to validate Crh deletion, and an Unpaired Student’s T-Test (p=0.0127) suggests there was a significant decrease in Crh punctae in cre treated CRF floxed mice when compared to control treated floxed mice, indicating that the Crh genetic deletion model works as intended (**supplemental Fig. S2**).

**Fig. 6.**
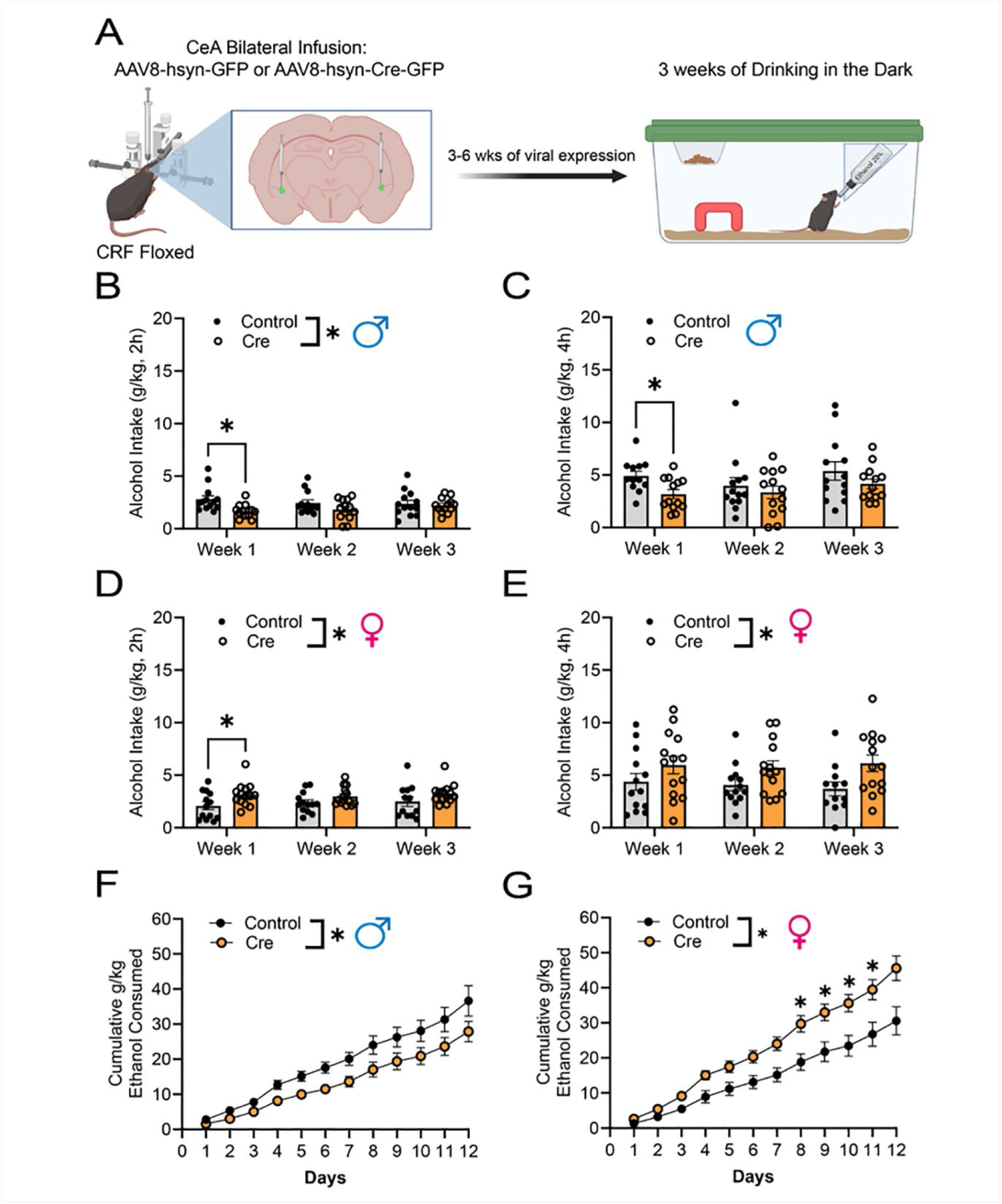
Genetic deletion of CRF in CeA neurons. (A) Experimental workflow beginning with bilateral AAV infusion to the CeA of CRF floxed mice, followed by 3-6 weeks of recuperation before engaging in 3 weeks of Drinking in the Dark (DID). Created using Biorender.com (B) CeA CRF genetic deletion in CRF floxed male mice decreased average weekly ethanol intake of the 2hr period in the first week of DID. (C) CeA CRF genetic deletion in CRF floxed male mice decreased average weekly ethanol intake of the 4hr period in the first week of DID (D) CeA CRF genetic deletion increased average weekly ethanol intake of the 2hr period in the first week of DID of the CRF floxed female mice. (E) CeA CRF genetic deletion had an overall increase on the weekly ethanol intake of the 4hr period in CRF floxed female mice despite each individual week showing no statistically significant differences. (F) CeA CRF genetic deletion reduced the cumulative intake of male CRF floxed mice despite no individual day indicating statistically reduced intake. (G) CeA CRF genetic deletion increased overall cumulative intake in female CRF floxed mice, with days 7-10 showing statistically significant differences when compared to control female mice. Closed circles denote AAV8-hsyn-GFP infused mice (Control), while open circles represent AAV8-hsyn-GFP-Cre infused mice (Cre), error bars are represented as SEM, *p < 0.05.

When comparing male Control and Cre treated mice in the weekly 2-hr consumption (**Fig. 6B**), mixed-effect analysis suggests that there was a main effect of viral treatment (F(1,24)=4.438, p=0.0458), but no main effect of time (F(2,47)=0.2971, p=0.7443) or time by viral treatment (F(2,47)=2.542, p=0.0895). Šídák multiple comparison post hoc tests between male control and cre treated CRF floxed mice indicates a significant decrease in 2hr weekly average alcohol consumption only at week 1 (p=0.0167) with no significant difference in week 2 (p=0.2945) or 3 (p=0.9622). When comparing the weekly 4hr alcohol intake of male control and cre treated CRF floxed mice (**Fig. 6C**), Mixed-effect analysis suggest there was only a main effect of time (F(1.654,38.88)=3.797, p=0.0386), with no main effect of viral treatment (F(1,24)=2.733, p=0.1113) or time by treatment (F(2,47)=0.9377, p=0.3987). Šídák multiple comparison post hoc tests between male Control and Cre treated CRF floxed mice indicates a significant decrease in 2hr weekly average alcohol consumption only at week 1 (p=0.0290) with no significant difference in week 2 (p=0.8855) or 3 (p=0.5381). Notably when comparing cumulative daily drinking of male Control and Cre treated CRF floxed mice (**Fig. 6F**), two-way repeated measures ANOVA suggest there was a main effect of time (F(1.177,25.89)=133.8, p<0.0001), viral treatment (F(1,22)=4.946, P=0.0367) and time by treatment (F(11,242)=2.031, p=0.0263). While there was an overall reduction in drinking in the male cre treated CRF floxed mice, Šídák multiple comparisons post-hoc tests indicate that this was not statistically significant at any of the 12 days. When comparing female Control and Cre treated CRF floxed mice 2hr weekly average alcohol consumption (**Fig. 6D**), mixed-effect analysis shows there was a main effect of viral treatment (F(1,25)=4.880, p=0.0366), but no main effect of time (F(1.821,44.62)=0.9119, p=0.4009) or time by treatment (F(2,49)=0.6031, p=0.5511). As opposed to male mice, Šídák multiple comparisons post-hoc tests indicate a significant increase in alcohol intake between Control and Cre treated female mice in week 1 (p=0.0328), but not week 2 (p=0.1085) and 3 (p=0.1551). Furthermore, when comparing female Control and Cre treated CRF floxed mice weekly 4hr alcohol intake (**Fig. 6E**), mixed-effect analysis show there was a main effect of viral treatment (F(1,25)=4.471, p=0.0446), and no main effect for time (F(1.862,45.61)=0.1569, p=0.8407) and time by viral treatment (F(2,49)=0.1903, p=0.8273). Despite the overall increase in alcohol intake in female Cre mice, Šídák multiple comparisons post-hoc tests indicate no significant differences between Control and Cre females in weeks 1 (p=0.4619), 2 (p=0.1874) or 3 (p=0.0772). Notably, when comparing cumulative daily drinking of female Control and Cre treated CRF floxed mice (**Fig. 6G**), two-way repeated measures ANOVA show there was a main effect in viral treatment (F(1,23)=9.228, p=0.0058), time F(1.212,27.88)=208.5, p<0.0001) and time by treatment (F(11,253)=7.396, p<0.0001). Šídák multiple comparison post-hoc tests indicate this increase in alcohol intake was statistically different in days 7 (p=0.0429), 8 (p=0.0393), 9 (p=0.0499), and 10 (p=0.0482). **Supplemental Fig.S3** illustrates the representative location where maximum virus was localized per animal for the Control and Cre CRF floxed male and female mice.

### CeA deletion of CRF1R

We wanted to determine if local CeA CRF1R plays a role in binge-like alcohol drinking, to accomplish this we used AAV delivery of Cre in combination with a CRF1R floxed mouse line to enable site specific deletion of the Crhr1 gene, and tested alcohol consumption during drinking in the dark (**Fig. 7A**). FISH was used to validate Crhr1 deletion, and an unpaired student’s T-test (p=0.0166) indicates that there was a significant decrease in Crhr1 punctae in Cre treated CRF1R floxed mice when compared to Control treated CRF1R floxed mice (**supplemental Fig. S4**). This indicates that the crhr1 genetic deletion model works as intended. When comparing both male and female Control and Cre treated CRF1R floxed mice, there was no main effect of viral treatment, time, and time by viral treatment at the 2hr weekly average alcohol consumption, 4hr weekly alcohol consumption, and cumulative alcohol consumption (**Fig. 7B-G**). This suggests that CRF1R synthesized in CeA neurons does not play a critical role in binge-like alcohol consumption. **Supplemental Fig. S5** illustrates the representative location where maximum virus was localized per animal for the Control and Cre CRF1R floxed male and female mice.

**Fig. 7.**
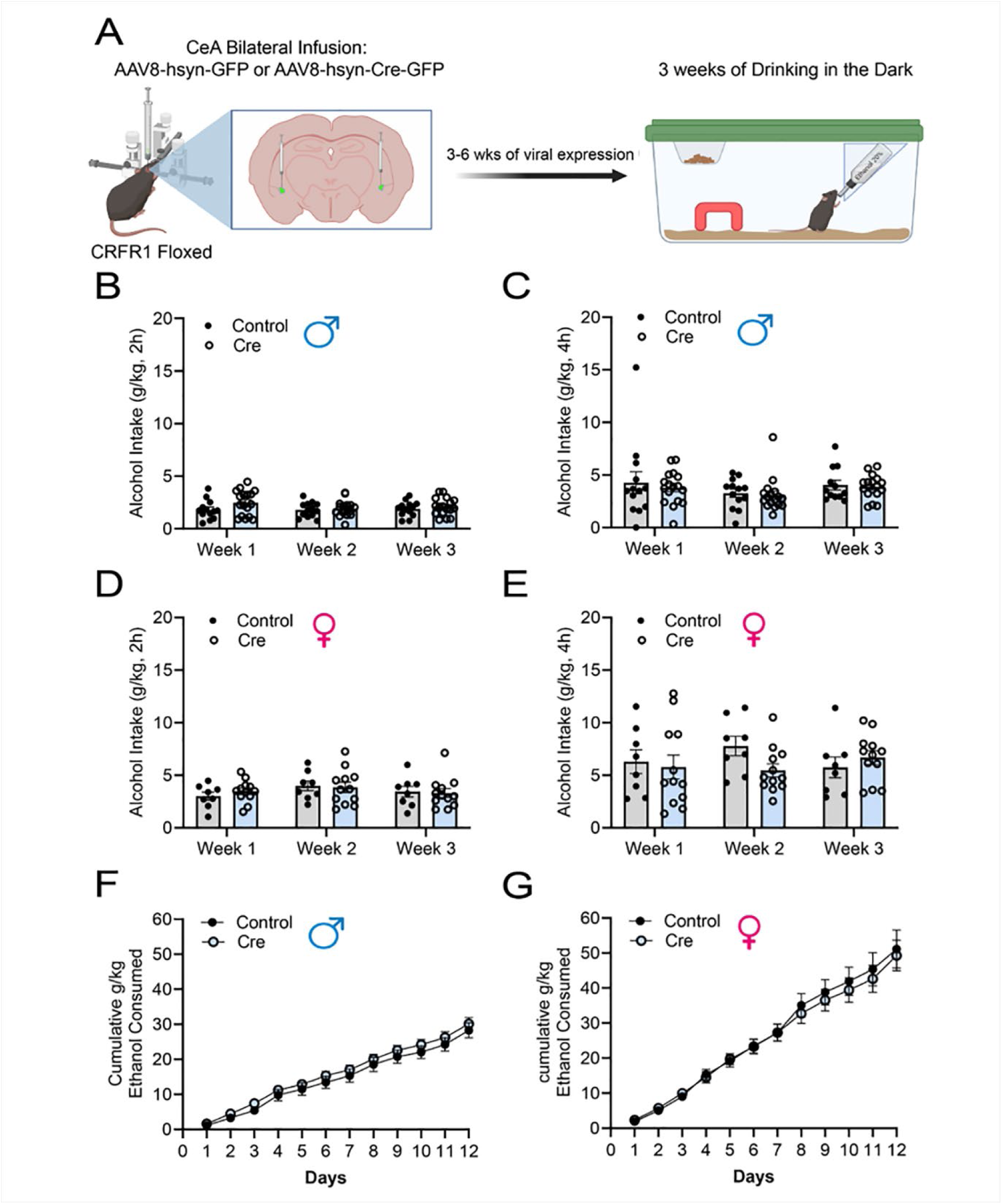
Genetic deletion of CRF1R on CeA neurons. (A) Experimental workflow beginning with bilateral AAV infusion to the CeA of CRFR1 floxed mice, followed by 3-6 weeks of recuperation before engaging in 3 weeks of DID.Created using Biorender.com (B) No effect of CeA CRFR1 genetic deletion in the average weekly ethanol intake of the 2hr period in CRFR1 flox male mice. (C) No effect of CeA CRFR1 genetic deletion in the weekly ethanol intake of the 4hr period in CRFR1 floxed male mice. (D) No effect of CeA CRFR1 genetic deletion in the average weekly ethanol intake of the 2hr period in CRFR1 floxed female mice. (E) No effect of CeA CRFR1 genetic deletion in the weekly ethanol intake of the 4hr period in CRFR1 floxed female mice. (F) No effect of CeA CRFR1 genetic deletion in the cumulative intake of male CRFR1 floxed mice (G) No effect of CeA CRFR1 genetic deletion in the cumulative intake of female CRFR1 floxed mice. Closed circles denote AAV8-hsyn-GFP infused mice (Control), while open circles represent AAV8-hsyn-GFP-Cre infused mice (Cre), error bars are represented as SEM.

## DISCUSSION

Our results here show that chemogenetically silencing a CRF+ circuit between the CeA and LH significantly blunts binge-like ethanol drinking in male, but not female, mice (**Fig. 1A**). This was associated with a significant reduction of BECs in male, but not female, mice (**Fig. 1B**) with no alterations in sucrose intake (**Fig. 1C**). Importantly, CNO treatment failed to alter ethanol intake, BECs or sucrose intake in mice treated with the control viral vector (**Fig. 2**). This CeA-LH CRF+ neurocircuitry has been mapped previously in rats and associated with avoidance of stress-associated stimuli^19^, and we provide evidence of a CRF+ CeA → LH circuit in our CRH-ires-cre mice here (**supplemental Fig. S6**). We extend the findings of Weera et al. by showing that the CRF+ CeA → LH neurocircuitry is not only involved in modulating stress, but also ethanol consumption, and that the contributions of these projections are male-specific. Further, the CeA → LH neurocircuitry we identified here acts as a specific neuronal target in binge-like ethanol consumption, since chemogenetic silencing of this pathway had no effect on sucrose consumption in either sex. And these effects were not a byproduct of CNO manipulation; by itself, CNO failed to cause any significant changes in ethanol consumption.

Pharmacological inhibition of CRF1R in the LH reciprocated these findings by blunting DID ethanol intake and associated BECs in male, but not female mice, but not altering sucrose intake (**Fig. 3**). Activation of the CRF2R did not significantly alter binge-like ethanol intake, BECs, or sucrose intake in either males or females (**Fig. 4**). Previous work in other brain regions has found pharmacological inhibition of CRF1R and activation of CRF2R both lead to a reduction in ethanol consumption^38,39^, however, our current results suggest that, in the LH, at the doses tested only the CRF1R is involved in modulating ethanol intake and only in male mice. Additionally, we found that a history of binge-like ethanol intake was associated with decreased CRF mRNA and elevated CRF2R mRNA in the CeA (**Fig. 5**). Finally, consistent with chemogenetic and pharmacological data, we found that genetic deletion of CRF from CeA neurons significantly blunted cumulative ethanol intake in male, but increased in female mice (**Fig. 6**), while genetic deletion of CRF1R from the CeA did not alter ethanol intake in either sex (**Fig. 7**). A schematic summarizing the central findings of our studies is presented in **Fig. 8**.

**Fig. 8.**
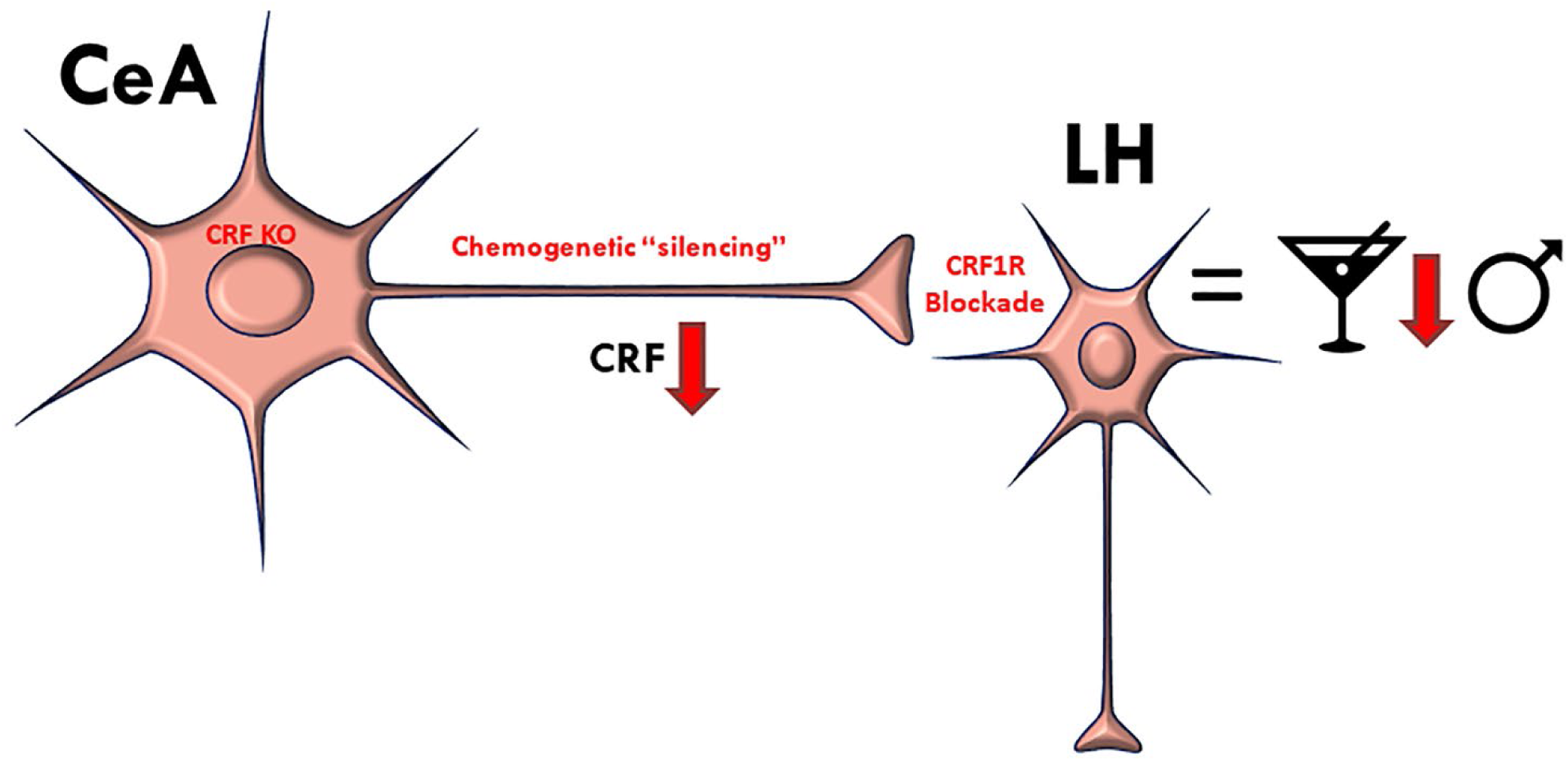
Schematic representation of central finding from chemogenetic, genetic deletion, and pharmacological studies. Chemogenetic “silencing of the CRF+ CeA ◊ LH circuit, genetic deletion of CRF in CeA neurons, and CRF1R blockade in the LH significantly blunted binge-like ethanol intake in male, but not female, mice.

CRF1R antagonism in the LH of male mice led to a reduction in binge-ethanol consumption, reflecting our male-specific findings in inhibiting CRF projections from the CeA to the LH; No such changes were observed using Ucn3 as a CRF2R agonist. Prior work in the CeA^38^, and the VTA^39^ have shown opposing effects of both CRF1R and CRF2R on binge-like ethanol consumption. While both the CeA and VTA are closely linked to the LH via CRF^40^, we were only able to observe the role of CRF1R antagonism in modulating ethanol intake in males. The sexually dimorphic role of CRF signaling, particularly via the CRF1R has been observed previously. As such, relative to males, female CRF mRNA, CRF1R binding, and the number of CRF+ neurons are upregulated through a number of brain regions such as the PVN^27,41–43^, amygdala^27,44^, and BNST^45,46^. At the level of cell signaling, acute application of ethanol onto mouse CeA slice preparations reduced GABA release onto CRF1R-expression neurons in male, but not female mice, and application of exogenous CRF increase the firing rate of CRF1R-expressing neurons to a greater extent in male mice^28^. When taken together, sex differences in CRF system organization and sensitivity may account for the observed sex differences in pharmacological and chemogenetic manipulations in the present report.

Future studies employing higher doses of CRF1R antagonists and/or CNO may reveal a role for the CRF+ CeA → LH circuitry in modulating ethanol intake in female mice which would support the hypothesis of sex differences in the sensitivity of this CRF circuit. Previously, females expressing Gi-DREADD in CRF+ BNST neurons required a higher dose of CNO to blunt ethanol binge drinking than males, suggesting that females are less sensitive to the effects of CNO on DREADD receptors^47^. Furthermore, intra-mPFC administration of CNO to activate Gi-DREADDS on NPY1R+ BLA terminals dose-dependently decreased ethanol intake in male and female animals, such that a low dose of CNO was effective at blunting ethanol consumption in males, while a high dose of CNO was required to achieve the same results in females^48^. In the current experiment, we did not assess different dosages of CNO. Thus, it is possible that our current findings of a male-specific CRF+ CeA → LH circuit in the modulation of binge-like ethanol intake might be recapitulated in females when using a higher dose of CNO. Likewise, we only tested one dose of the CRF1R antagonist NBI-35965 and CRF2R agonist Ucn3, and with sex differences in amygdala CRF1 binding^44^, it is possible that running a dose-response curve is necessary to capture the role of LH CRF1R and/or CRF2R in modulating females binge-like ethanol intake.

It is unlikely LH CRF1R signaling modulates binge-like ethanol intake via a mechanism that involves the hypothalamic-pituitary-adrenal axis (HPA) functioning as binge-like ethanol intake was not found to alter HPA-axis activity^49^. Consistently, repeated cycles of binge-like ethanol intake did not alter CRF immunoreactivity in the paraventicular nucleus of the hypothalamus, but did increase CRF immunoreactivity in the CeA, and this change lasted 18-24h into abstinence in male mice^13^. These previous observations and the current data suggest that while hypothalamic CRF1R signaling modulates binge-like ethanol intake, this signaling likely stems from extrahypothalamic sources of CRF (i.e., from the CeA) and does not involve hypothalamic pools of CRF that are involved with HPA-axis function.

We also found that repeated cycles of binge-like ethanol consumption altered CRF and CRF2R mRNA expression in the amygdala, but not amygdala CRF1R mRNA expression. Mice that received three cycles of DID showed significantly reduced CRF mRNA expression in the amygdala compared to water drinking control mice, while mice that received three or six cycles of DID showed significantly greater CRF2R mRNA expression in the amygdala compared to water drinking control mice. CRF2R expression remained significantly elevated 24 hours into abstinence. Previous research has demonstrated that CRF protein expression in the amygdala is elevated following binge-like ethanol consumption^13^, and CRF mRNA expression has been found to increase following ethanol dependence in rats^50^. We hypothesize that our data provide evidence for a compensatory mechanism where a decrease in CRF mRNA in the amygdala following binge-like ethanol consumption may serve as an attempt to attenuate increased CRF signaling and protein levels following ethanol intake, and failure to regulate this CRF system may contribute to the development of ethanol dependence, supporting an allostatic view of neurobiological changes in response to ethanol^51^. Neither CRF1R nor CRF2R mRNA expression in the LH were impacted by binge-like ethanol consumption. One limitation of the current data set is that, due to the small size of these brain regions in mice, two tissue samples were pooled to obtain sufficient mRNA for cDNA synthesis, effectively halving our sample size. Therefore, it is likely that the current qPCR study was underpowered to detect sex differences in mRNA expression. However, an inspection of the individual data points from the mRNA data (**Fig. 5**) does not suggest any obvious trends for sex differences in expression patterns.

One of the key questions that can arise through the use of DREADD approaches is that what is the cellular effector driving the change in behavior. For our work, one important question was if CRF from the CeA was driving these behavioral changes. While our pharmacology studies provided strong support of this idea, the genetic approach with site specific deletion could add converging evidence. In keeping with the pharmacology, we found that deletion of CRF in the CeA reduced ethanol consumption in male mice, yet increased consumption in female mice. The increase in ethanol drinking in female mice is especially intriguing when considering our DREADD findings, as it suggests an important sex difference in the modulatory systems and circuits that regulate behavior. In contrast, the deletion of CRF1R in the CeA had no effect on ethanol consumption. This is in opposition to the pharmacological studies supporting a role for CRF1R in the CeA in alcohol drinking. However, it is important to note that we deleted receptors in CeA neurons, leaving receptors on terminals from other regions intact, pointing to potential input regions as key drivers of the CRF1R driven pharmacological effects. It is also possible that the level of deletion that we found was insufficient to drive changes in function. Another limitation to our study is that virus spread leaked into other neighboring regions of the CeA including the BLA, which means we cannot rule out possible unwanted deletion of CRF or CRF1R in those regions causing non-specific effects.

In conclusion, the present observations reveal a novel CRF+ circuit between the CeA and LH that modulates binge-like ethanol intake in male, but not female mice. Given the abundance of research demonstrating that female rodents consume significantly more ethanol than male rodents, the present findings may represent a critical circuit that contributes to sex differences in ethanol intake. Theoretically, blunted sensitivity of this circuit in female mice may contribute to their elevated levels of ethanol intake. The present observations may also provide insight into sex-dependent treatment approaches for treating AUDs.

## Acknowledgments

Special thanks to Rhiannon Thomas, MS, for technical assistance with this project. This work was supported by NIH grants AA022048, AA013573, AA025809, F31AA029935, and T32DA007244. Partly funded by the HHMI James H. Gilliam Jr. Fellowship for Advanced Study (GT13514)

## Supplemental Materials

**Inhibiting CRF Projections from the Central Amygdala to Lateral Hypothalamus and Amygdala Deletion of CRF Alters Binge-Like Ethanol Drinking in a Sex-Dependent Manner**

### Supplemental Figures

**Fig. S1:**
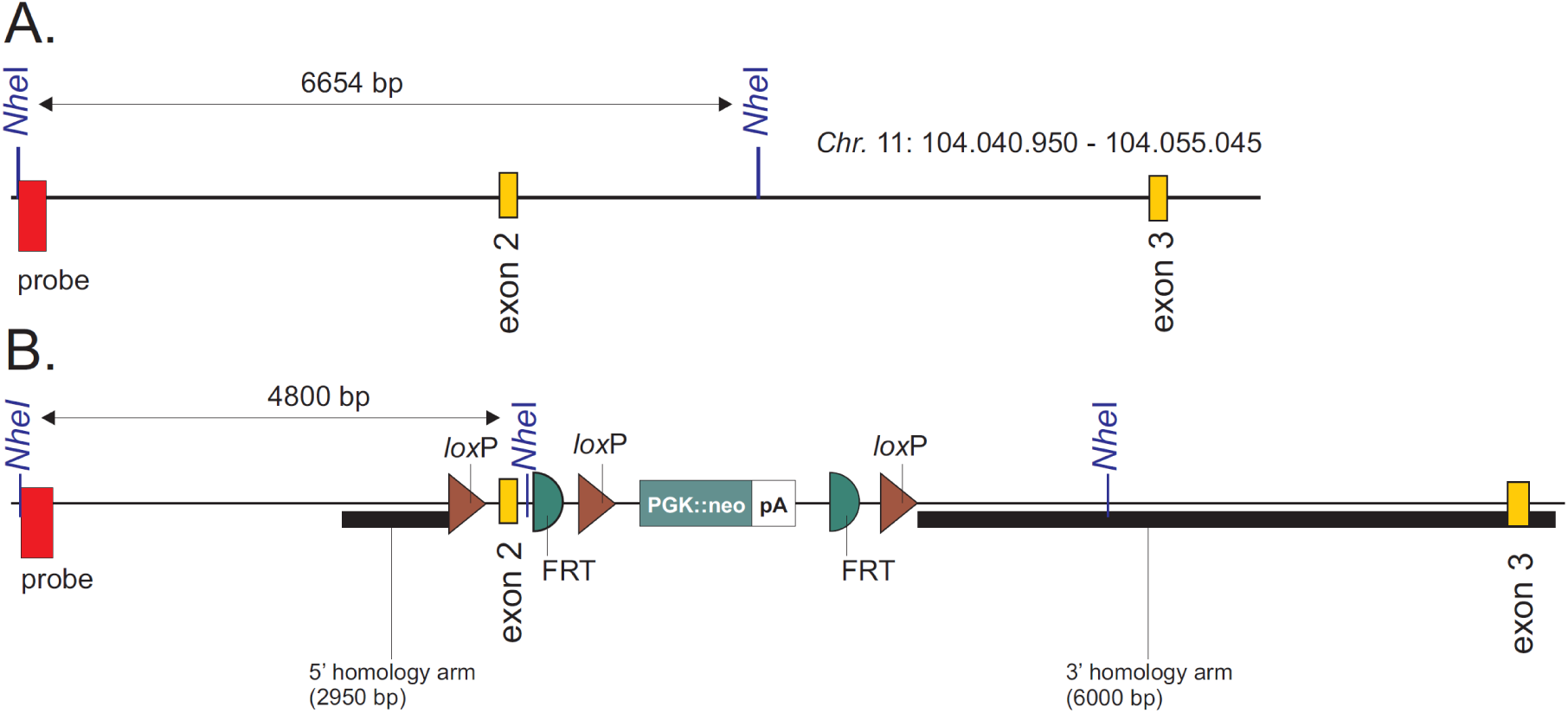
Targeting construct for generation of floxed CRFR1 mice.

**Fig. S2:**
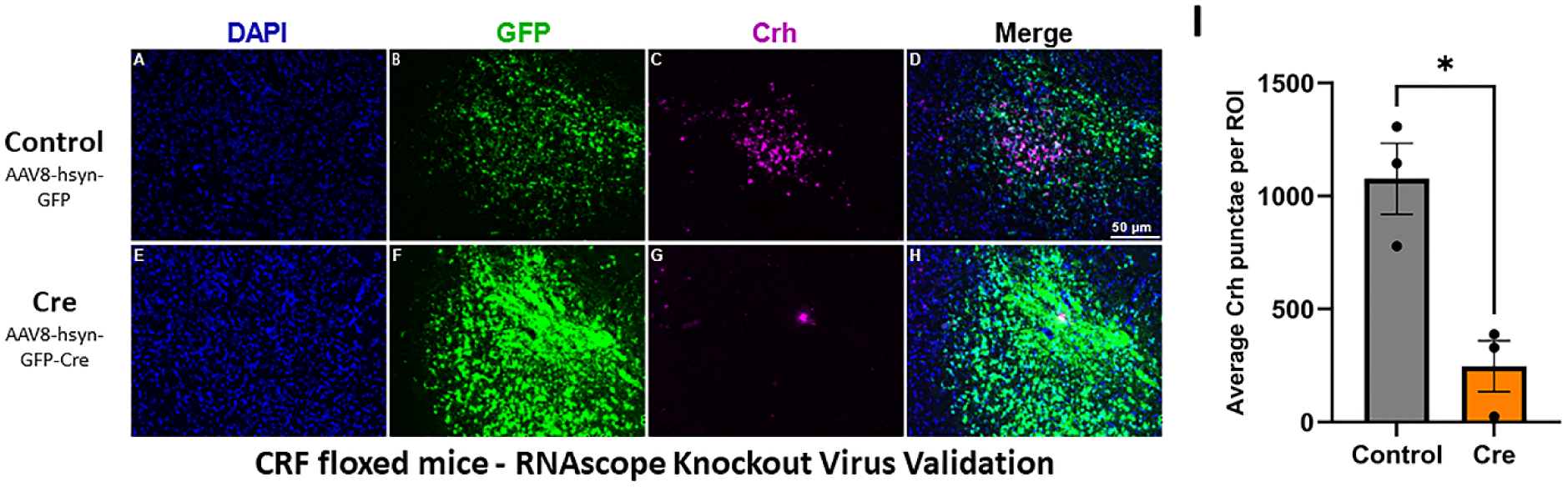
Validation of CRF genetic deletion. (A-D) Representative image showing Crh expression in the CeA of control treated mice (E-H) Representative image showing Crh expression in the CeA of Cre treated mice (I) Significant reduction of average Crh punctae per ROI in Cre treated mice. AAV8-hsyn-GFP treated CRF floxed mice (Control), AAV8-hsyn-GFP-Cre treated CRF floxed mice (Cre), scalebar 50um, each data point represents average punctae per ROI from each mouse, Control treated group had, N=3 mice, 4-10 serial sections per mouse, while Cre treated group had, N=3 mice, 2-5 serial sections per mouse. Crh was only counted in ROI’s that had GFP expression indicative of virus presence. *p < 0.05. DAPI, denotes nucleus of cells in blue, GFP denotes viral GFP tag in green, Crh in magenta denotes mRNA punctae of Crh. Error bars are represented as SEM, *p < 0.05.

**Fig. S3:**
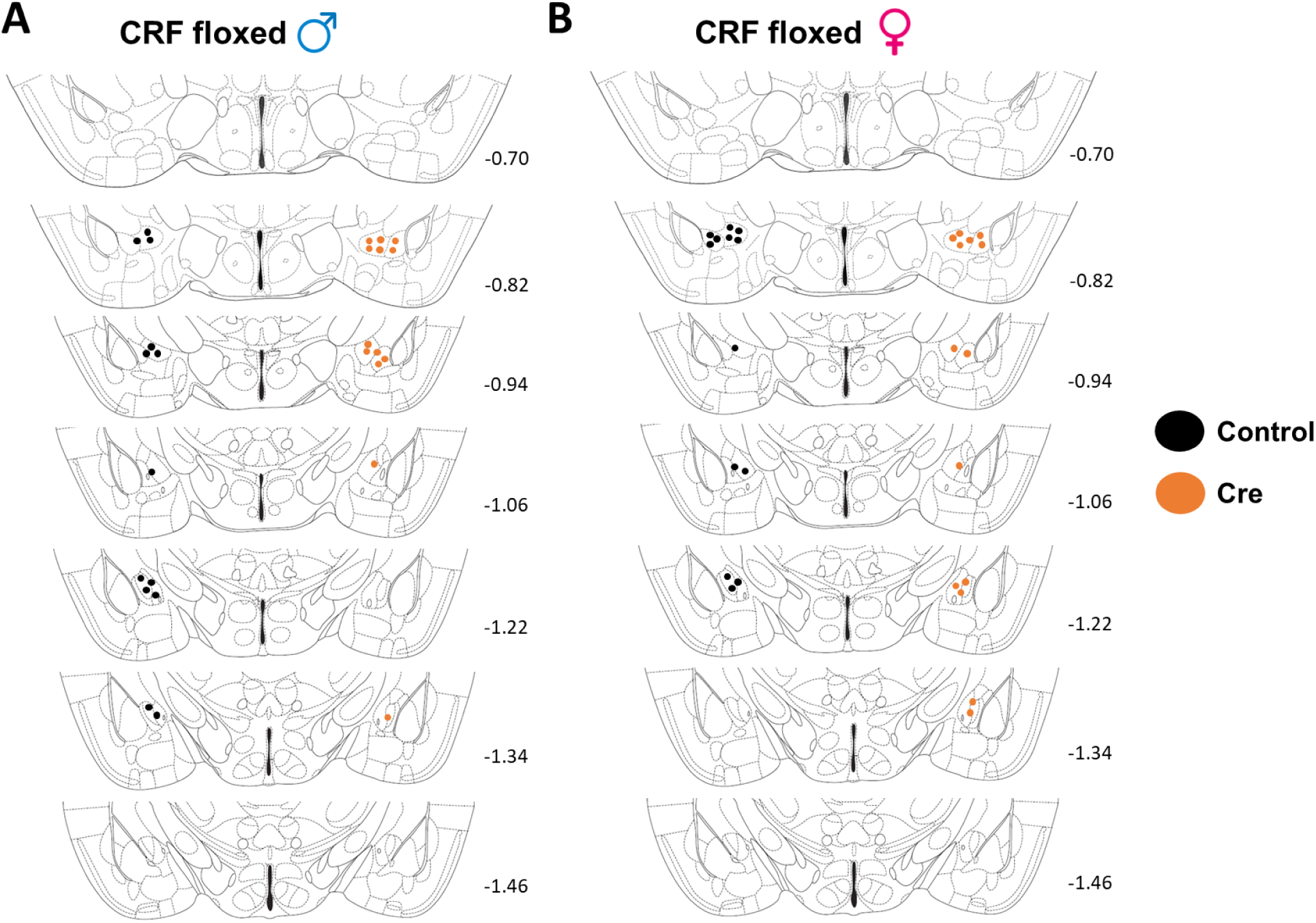
(A) Representative location relative to bregma where maximum virus was localized in male CRF floxed mice. (B) Representative location relative to bregma where maximum virus was localized in female CRF floxed mice. 1 dot per mouse, black dots = Control mice and orange dots = Cre mice.

**Fig. S4:**
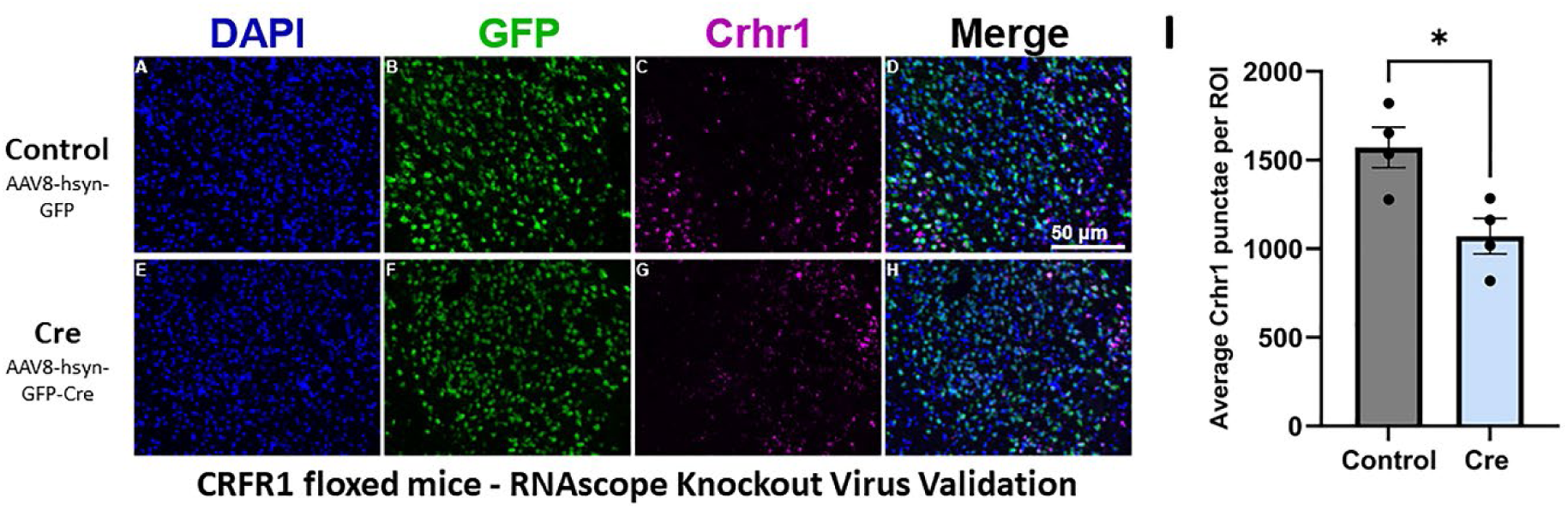
Validation of CRF1R genetic deletion. (A-D) Representative image showing Crhr1 expression in the CeA of control treated CRFR1 floxed mice (E-H) Representative image showing Crhr1 expression in the CeA of Cre treated CRFR1 floxed mice (I) Significant reduction of average Crhr1 punctae per ROI in Cre treated CRFR1 floxed mice. AAV8-hsyn-GFP treated CRFR1 floxed mice (Control), AAV8-hsyn-GFP-Cre treated CRFR1 floxed mice (Cre), scalebar 50um, each data point represents average punctae per ROI in each mouse, N=4 mice, 5-9 serial sections per mouse, while Cre treated group had, N=4 mice, 4-9 serial sections per mouse. Crhr1 was only counted in ROI’s that had GFP expression indicative of virus presence. Crhr1 was only counted in ROI’s that had GFP expression indicative of virus presence. *p < 0.05. DAPI, denotes nucleus of cells in blue, GFP denotes viral GFP tag in green, Crhr1 in pink denotes mRNA punctae of Crh. Error bars are represented as SEM

**Fig. S5:**
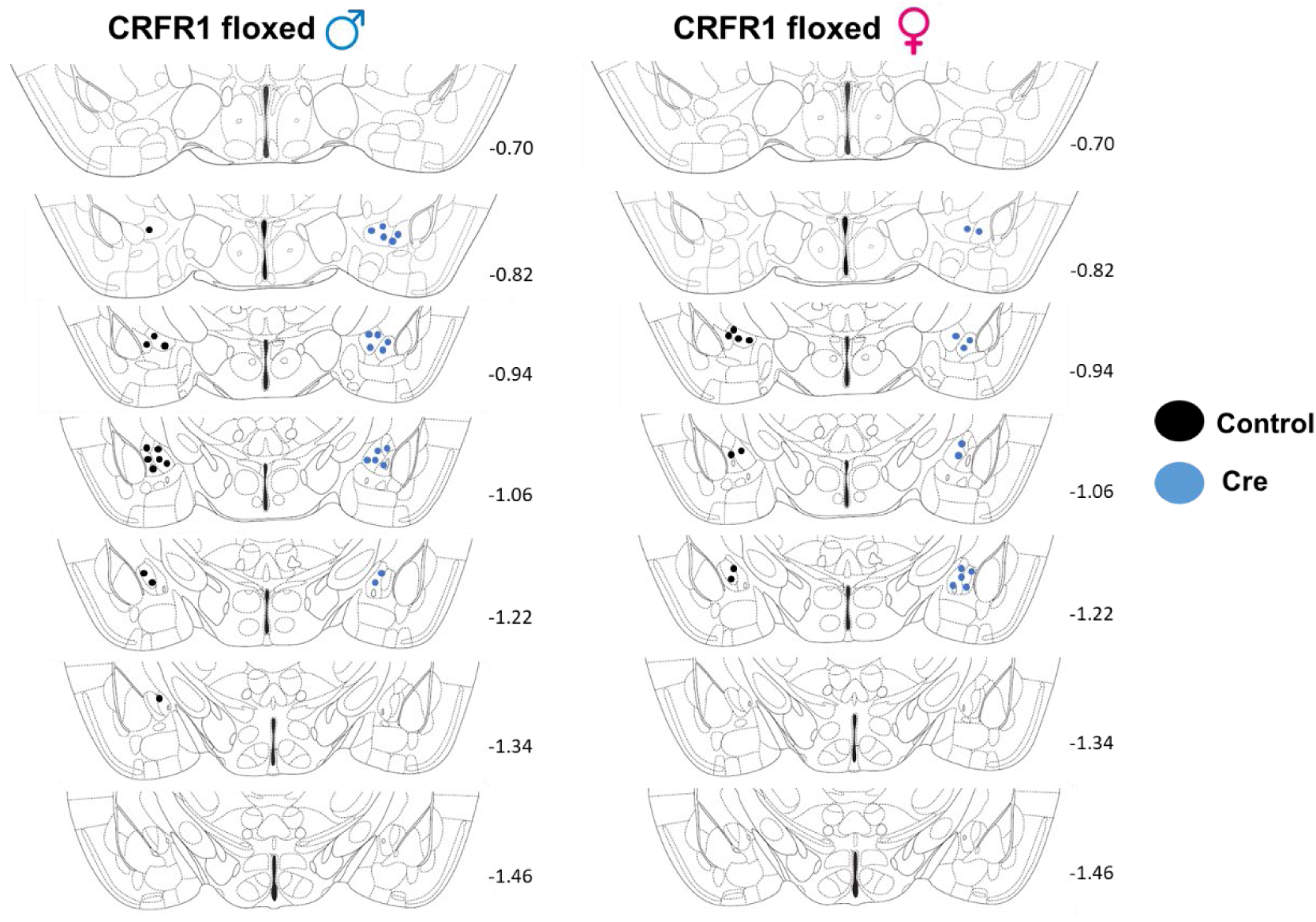
(A) Representative location relative to bregma where maximum virus was localized in male CRFR1 floxed mice. (B) Representative location relative to bregma where maximum virus was localized in female CRFR1 floxed mice. 1 dot per mouse, black dots = Control mice and blue dots = Cre mice.

**Fig. S6:**
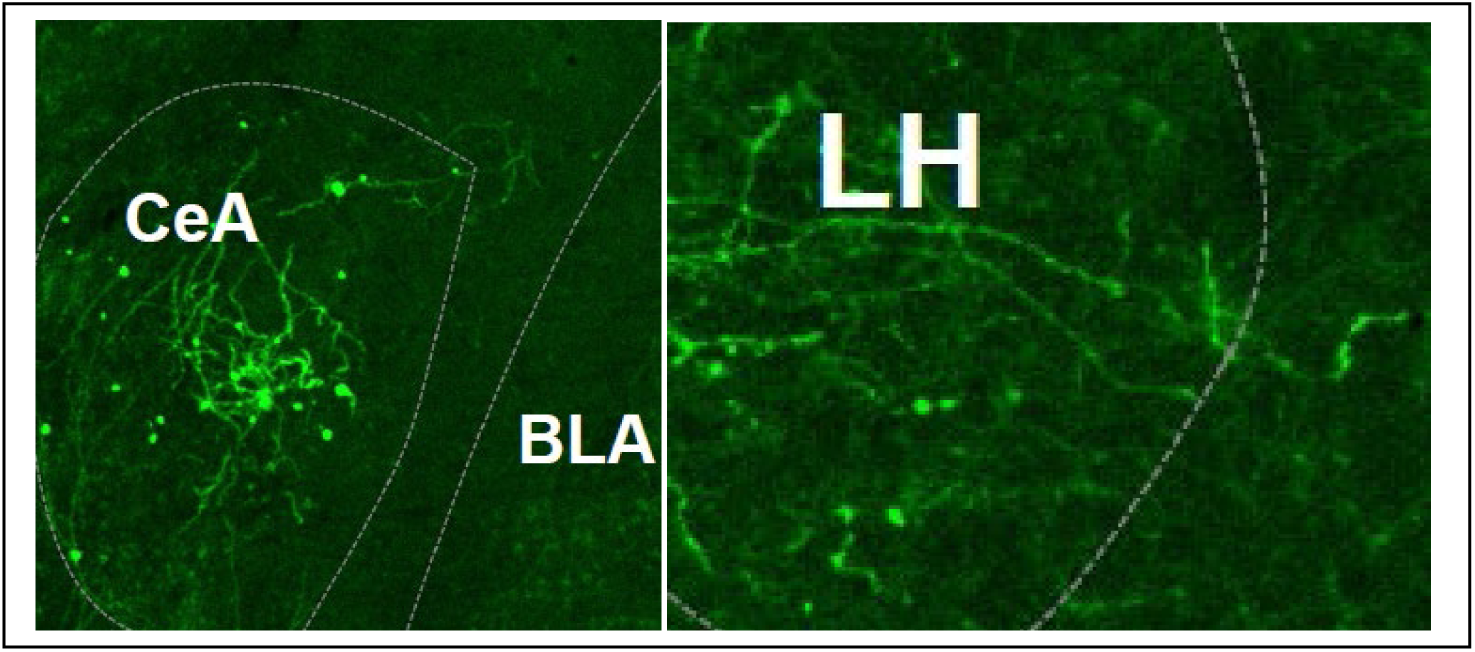
CRH-ires-cre mice were given of a cre-inducible reporter virus (AAV5-EF1a-DIO-hChR2-mCherry; 0.3 µl bilaterally) into the central amygdala (CeA) and 3-weeks later slices were collected containing the CeA (and basolateral amygdala; BLA) as well as the lateral hypothalamus (LH). Soma of CRF+ neurons are evident in the CeA (left panel) and CRF+ terminals a are evident in the LH (right panel).

### Extended Methods

#### “Drinking in the Dark” (DID) Procedures

This procedure involves replacing the cage water bottle with a 20% (v/v) ethanol solution or a 3% (w/v) sucrose solution 3-hours into the animals’ dark cycle. Drinking access each day was limited to 2 hours, with ethanol consumption recorded at the end of each access session. The amount consumed by each subject was recorded during hour 1 and hour 2 on the test (4th) day. To correct for spillage, in parallel to each experiment we ran a drip bottle, 20% ethanol bottle in an empty cage. Sucrose drinking control animals went through the same DID procedure with a 3% (w/v) sucrose solution, and consumption was recorded at the end of each access session (with 1-and 2-hour consumption levels recorded on the last day). Animals received a 3-day period of abstinence before the new 4-day cycle of DID began. For the genetic deletion experiments, alcohol drinking session consisted of 2 hours during the first three days (Monday-Wednesday) and to 4hrs (Thursday) during the 4^th^ day of each week for 3 weeks, alcohol bottles were weighted after each drinking session each day. Mouse weights were measured at least once a week for the duration of the experiment. There were instances where a mouse weight was accidentally not recorded, in which case, the estimated weight using the calculated average weight-gain was used. Additionally, there were instances where the drip bottles leaked excessively. In these circumstances, the average drip value of the two other weeks was used. For the qPCR experiment, mice were randomly assigned to one of four groups while ensuring equal distribution of sex and initial body weight. Group one served as the water control group and remained ethanol-naïve for the duration of the experiment. Group two and three received three and six, 4-day cycles of DID, respectively, and brains were extracted immediately following the final session of DID. Group four received six cycles of DID followed by a 24 hour period of abstinence from ethanol, therefore brains were extracted 24 hours following the final session of DID.

#### Generation of Floxed CRFR1 Mice

A phage DNA library was screened in two rounds with a 32P-dCTP-labeled probe corresponding to exon 2 (88 bp) of the Crhr1 gene (MGI: 88498) and identify mouse genomic DNA clones that contains exon 2. Several clones were picked and mapped with a panel of restriction enzymes. Clone 12 was selected and used to create the targeting construct. The construct included 1 kb genomic sequence as 5’ arm of recombination, a first loxP site, ∼750 bp genomic sequence containing exon 2, a pair of neighboring frt and loxP sites, a pGK1-neo-pA resistance gene cassette, a second pair of frt and loxP sites, 6 kb genomic sequence as 3’ arm of recombination (see Figure), and the herpes simplex virus thymidine kinase (TK) for negative selection. TL-1 embryonic stem (ES) cells derived from 129/SvEvTac mice cells were electroporated with the linearized vector and grown on fibroblast feeder cells in DMEM supplemented with 15% fetal bovine serum, 50 mg/ml gentamicin, 1000 U/ml leukemia inhibitory factor, 90 mM b-mercaptoethanol, 0.2 mg/ml G418 (positive selection), and 2 mM ganciclovir (negative selection). Three hundred and seventy-one independent neomycin-resistant colonies were selected and grown in 96-well plates on feeder layer, expanded, and analyzed for the presence of the mutant gene by performing Southern blot analysis using genomic DNA digested with NheI and hybridized with a 32P-labeled probe consisting of 304 bp upstream of the 5’ arm of recombination. Five positive clones (clones showing a 4.8 kB NheI fragment) were selected and screened at the 3’ end. Clones 3C6 and 4C4 were confirmed and injected into C57BL/6J blastocysts generating 4 and 3 chimeric animals (>60% brown fur), respectively. After demonstrating germline transmission for line 4C4, the mice were crossed to E2a-CRE mice to obtain partial recombination, i.e. 3 loxP to 2 loxP with the loss of the neomycin-resistance gene cassette. The mice were then bred for additional generations into C57BL6/J prior to study. The targeting construct for generation of floxed CRFR1 mice is presented in **supplemental Fig S1**.

#### Fluorescence In Situ Hybridization (FISH)

In order to confirm genetic deletion of CRF and CRFR1, brain collection, cryosectioning and FISH was performed as in (Bloodgood et al, 2021). Mouse brains were flash frozen in 2-methylbutane and dry ice and then stored in −80°C. 14µm-thick serial sections were collected using a Leica CM 3050S cryostat (Leica Biosystems), these sections were mounted on Super Frost Plus Slides (Fisher Scientific), 4-5 sections per slide. FISH was performed following the manufacture’s protocol for the Advanced Cell Diagnostics RNAscope Fluorescence Multiplex Assay (Advanced Cell Diagnostics). We used the following probes for detecting mRNA puncta, Crh-C1 (Mm-Crh), Crhr1-C2 (Mm-Crhr1-C2), eGFP-C3 (EGFP-O4-C3). In order to fluorescently view the mRNA puncta we used TSA vivid Dyes, opal 570 or opal 650. The nucleus of the cells were counterstained with RNAscope DAPI for visualization. Finally, cover slips were mounted on the slides using ProLong Diamond Antifade Mountant (Thermo Fisher Scientific). Slides were imaged using a VS200 Slide Scanner microscope. Images were analyzed using QuPath software version 0.5.0, and an ROI was created over the CeA. In the ROI, the cell detection function (using a value of a 100-intensity parameter threshold) was used to detect all DAPI-positive cells while the subcellular spot detection function (using a detection threshold of 250), was used to detect Crh or Crhr1 puncta, in DAPI-positive cells. Only ROIs that had GFP, indicating the presence of virus, were included for downstream analysis in this study.

#### Perfusion and Histology

For cannula placement checks and DREADD expression, mice were prepared for perfusion by administering a 0.1mL/kg i.p. injection of ketamine/xylazine (6.67 mg/0.1 mL; 0.67 mg/0.1 mL; in 0.9% saline). Then, mice were perfused transcardially with 0.1M phosphate buffer saline (PBS; pH=7.4) and10% buffered formalin phosphate (fisher chemical). After extraction, brains were fixed in 10% buffered formalin phosphate for 24-48 hours, before being sliced at 40µm thickness on a vibratome (Leica VT1000S vibratome; Wetzlar, Germany). For the chemogenetic study, cannula placement and DREADD expression were verified on an optical wide field microscope (Leica DM6000) with a digital camera (Roper Scientific). Cannula placement checks for the pharmacological experiments were conducted using a Nikon e400 biological microscope with a digital sight ds-u1 imaging attachment (Nikon Instruments Inc., Melville, NY, USA). Cannula targets were determined by the end of the guide cannula plus 2mm to account for the injector tip. For the genetic deletion experiments, to assess targeting of AAV in floxed mouse lines, tissue was prepared as before (Bloodgood et al., 2021). Mice were sacrificed one day after completing 3 weeks of DID. Mice were perfused using PBS and 4% PFA. Brains were isolated and stored in 4% PFA and later washed and stored in PBS at 4°C. Brain slices were serially sectioned at 45µm-thick using a vibratome (VT 1200s, Leica Biosystems), and sections were collected in Super Frost Slides and mounted using Vectashield Hardset Antifade mounting media with DAPI (Vector Laboratories). To assess targeting of virus, brain sections were then imaged using a VS200 Slide Scanner (Olympus) with Orca Fusion camera (Hamamatsu). Mice that had bilateral and unilateral hits were included in the genetic deletion study.

#### Reverse Transcription Quantitative Polymerase Chain Reaction (RT-qPCR)

For mRNA expression analysis, mice were euthanized via rapid decapitation and trunk blood samples were collected for BEC analysis. Brain tissue was extracted and flash frozen in O.C.T. compound (ThermoFisher Scientific, Waltham, MA). Brains were sliced at 40µm thickness on a cryostat (CM3050 S, Leica, Buffalo Grove, IL) and 1mm tissue punches were collected bilaterally for the amygdala and LH. Tissue samples from two animals within the same group and sex were pooled for each sample to obtain sufficient RNA for analysis. Tissue punches were collected in bead homogenizing tubes containing 500 mL of TriReagent (Molecular Research Center, Cincinnati, OH). RNA extraction, cDNA synthesis, and RT-qPCR were performed as previously described (Barkell et al., 2022; Paniccia et al., 2018). Briefly, RNA from each sample was extracted and purified using the Qiagen RNEasy Tissue Mini Kit for RNA purification (Qiagen, Hilden, Germany). RNA spectroscopy using the Take3 Application and Gen5 Software for Nucleic Acid Quantification (BioTek Instruments Inc., Winooski, VT) was used to assess the concentration and purity of the RNA in each sample. Each sample was diluted with PCR grade water so that all samples contained an equal concentration of RNA. cDNA was synthesized using the Advantage for RT-PCR Kit (Clontech, Takara, Mountain View, CA) and a Veriti 96 Well Fast Thermal Cycler (Applied Biosystems, ThermoFisher Scientific, Waltham, MA). qPCR was performed using the TaqMan Fast Advanced Master Mix Kit (Applied Biosystems, ThermoFisher Scientific, Waltham, MA) and TaqMan Gene Expression Assays for CRF (Mm04206019_m1), CRF1R (Mm00432670_m1), CRF2R (Mm00438308_m1), and the housekeeping gene BetaActin (Mm01205647_g1). Samples underwent repeated cycles of amplification and data collection using the QuantStudio 6 Flex real-time PCR system (ThermoFisher Scientific, Waltham, MA). Additionally, cDNA samples from amygdala and LH tissue were sent to the UNC Chapel Hill Advanced Analytics Core for further qPCR analysis.

#### Animal exclusion criteria

Animals were removed from analysis if they were found to be a significant outlier via the Grubbs test (Alpha=0.05), if a cannula misplacement (unilateral or bilateral) was found, DREADD expression was misplaced/missing, if GFP expression (as a marker of cre deletion) was misplaced or missing, or if they drank less than 0.30 g /kg in the first week. In total, 2 females and 3 males from the Gi DREADD group, and 7 females and 4 males from the control DREADD group were excluded due to no or unilateral DREADD expression. Additionally, 1 female receiving 6 cycles of DID was excluded from analysis of CRF mRNA expression in the amygdala and 1 female and 1 male receiving 3 cycles of DID were excluded from analysis of CRF1R mRNA expression in the amygdala due to significantly outlying data. No animals from the pharmacology experiment had misplaced cannula.

